# Estimation of the number of synapses in the hippocampus and brain-wide by volume electron microscopy and genetic labeling

**DOI:** 10.1101/2020.02.18.953802

**Authors:** Andrea Santuy, Laura Tomás-Roca, José-Rodrigo Rodríguez, Juncal González-Soriano, Fei Zhu, Zhen Qiu, Seth GN Grant, Javier DeFelipe, Angel Merchan-Perez

## Abstract

Determining the number of synapses that are present in different brain regions is crucial to understand brain connectivity as a whole. Membrane-associated guanylate kinases (MAGUKs) are a family of scaffolding proteins that are expressed in excitatory glutamatergic synapses. We used genetic labeling of two of these proteins (PSD95 and SAP102), and Spinning Disc confocal Microscopy (SDM), to estimate the number of fluorescent puncta in the CA1 area of the hippocampus. We also used FIB-SEM, a three-dimensional electron microscopy technique, to calculate the actual numbers of synapses in the same area. We then estimated the ratio between the three-dimensional densities obtained with FIB-SEM (synapses/μm^3^) and the bi-dimensional densities obtained with SDM (puncta/100 μm^2^). Given that it is impractical to use FIB-SEM brain-wide, we used previously available SDM data from other brain regions and we applied this ratio as a conversion factor to estimate the minimum density of synapses in those regions. We found the highest densities of synapses in the isocortex, olfactory areas, hippocampal formation and cortical subplate. Low densities were found in the pallidum, hypothalamus, brainstem and cerebellum. Finally, the striatum and thalamus showed a wide range of synapse densities.

## Introduction

Determining the number of synapses that are present in different brain regions is crucial to understand brain connectivity as a whole. Synapses can be identified with several methods, including genetic labeling of synaptic scaffolding proteins and electron microscopy (EM). Membrane-associated guanylate kinases (MAGUKs) are a family of scaffolding proteins that participate in the regulation of cell polarity, cell adhesion and synaptic signal transduction (Migaud et al., 1998; Ye et al., 2018; Zhu et al., 2016). PSD95 and SAP102 belong to the MAGUK family and are expressed in the postsynaptic density (PSD) of excitatory glutamatergic synapses (Aoki et al., 2001; Chen et al., 2018, 2008; DeGiorgis et al., 2006; Farley et al., 2015; Husi et al., 2000; Petersen et al., 2003; Valtschanoff et al., 1999; Yamasaki et al., 2016), where they contribute to the recruitment and retention of glutamate receptors (Hafner et al., 2015; Jeyifous et al., 2016; Levy et al., 2015). Genetic labeling of the endogenous PSD95 and SAP102 postsynaptic proteins and imaging using Spinning Disk confocal Microsocpy (SDM) have been proven to be useful for the characterization of synapse diversity in all brain regions of the mouse. SDM is a rapid method that allows the imaging of entire brain sections, so the simultaneous visualization of millions of synapses is made possible, obtaining bi-dimensional densities of fluorescent puncta per surface area (puncta/100 μm^2^) (Zhu et al., 2018).

Previous attempts have been made to calculate the density of synapses in the brain using EM. This technique allows the identification of individual synapses, although it is restricted to much smaller fields of view. Furthermore, most of these EM studies apply stereological techniques to a limited number of EM sections. Although stereology is a proven valuable method for object counting, the total number of synapses is an estimation which is subject to several technical limitations [see (DeFelipe et al., 1999) for a review]. In the present study, we use Focused Ion Beam milling-Scanning Electron Microscopy (FIB-SEM). With this technique, sectioning and imaging are fully automated, allowing the acquisition of multiple serial micrographs. Later, the micrographs can be stacked with the help of software tools, such that they represent a three-dimensional sample of tissue (Merchán-Pérez et al., 2009). In this way, all individual synapses can be identified and counted within a known volume of brain tissue, and thus the true density of synapses per unit volume can be obtained directly (not through estimations using stereological methods).

The aim of our study was twofold. First, we wanted to obtain detailed data about the density and size of synapses in the hippocampus. To this end, we used SDM to measure the densities of PSD95 and SAP102 puncta in stratum oriens (SO), stratum radiatum (SR) and stratum lacunosum-moleculare (SLM) of CA1. We also used FIB-SEM to measure the actual density of synapses in three-dimensional samples of the same strata (Figure 1). We then calculated the quantitative relationship between the densities and sizes of fluorescent puncta and synapses obtained by the two methods. Second, given that volume electron microscopy cannot be applied brain-wide, we wanted to obtain an estimate of the number of synapses in other regions of the brain where measurements of PSD95 and SAP102 puncta were available (Zhu et al., 2018). We based this estimate on the quantitative relationship or conversion factor between SDM and FIB-SEM data previously obtained in the hippocampus. Even though this approach has several limitations and underestimates the actual numbers of synapses, it provides valuable information on the *minimum* number of excitatory synapses that are present in more than a hundred brain regions.

**Figure 1.**
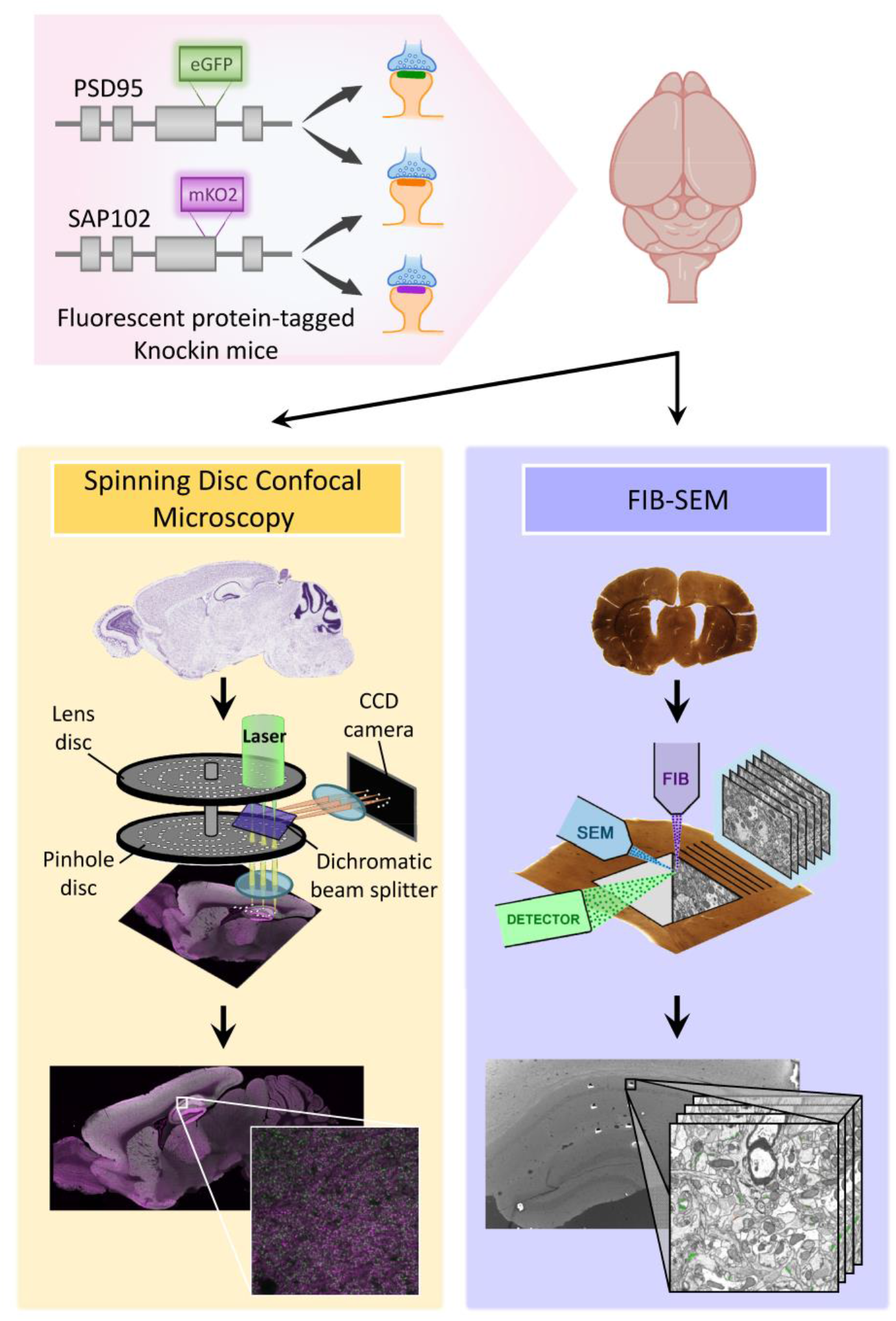
General methodology. Knockin mice expressing fluorescent PSD95 and SAP102 were imaged with Spinning Disc Confocal Microscopy (SDM) and with FIB-SEM. SDM allows the acquisition of large field, 2D fluorescent images, while FIB-SEM is an electron microscopy technique with a resolution in the scale of nanometres that generates 3D stacks of images, but with a smaller field of view. We have used a combination of both techniques to estimate the actual densities of synapses per unit volume brain-wide.

## Materials and Methods

### Animals

For this study, we used adult male mice (postnatal day 56) expressing fluorescently labeled PSD95 and SAP102 postsynaptic proteins (PSD95^eGFP/eGFP^; SAP102^mKO2/Y^) (Zhu et al., 2018). All animals were handled in accordance with the guidelines for animal research set out in the European Community Directive 2010/63/EU, and all procedures were approved by the Ethics Committee for Animal Experimentation of the Cajal Institute (CSIC, Spain).

### Tissue preparation for spinning disc microscopy

Sixteen mice were anesthetized by an intraperitoneal injection of 0.1 mL of 20% w/v sodium pentobarbital (Euthatal, Merial Animal Health Ltd. or Pentoject, Animalcare Ltd.). After complete anesthesia, 10 mL of phosphate buffered saline (PBS; Oxoid) were perfused transcardially, followed by 10 mL of 4% v/v paraformaldehyde (PFA; Alfa Aesar). Whole brains were dissected out and post-fixed for 3–4 h at 4° C in 4% PFA, and then cryoprotected for 3 days at 4 °C in 30% sucrose solution (w/v in 1× PBS; VWR Chemicals). Brains were then embedded into optimal cutting temperature (OCT) medium within a cryomould and frozen by placing the mould in isopentane cooled down with liquid nitrogen. Brains were then sectioned, with a thickness of 18 μm, using an NX70 Thermo Fisher cryostat, and cryosections were mounted on Superfrost Plus glass slides (Thermo scientific) and stored at −80 °C.

### Histology and immunohistochemistry

Sections were washed for 5 min in PBS, incubated for 15 min in 1 μg/mL DAPI (Sigma), washed and mounted using home-made MOWIOL (Calbiochem) containing 2.5% anti-fading agent DABCO (Sigma-Aldrich), covered with a coverslip (thickness #1.5, VWR international) and imaged the following day.

### Spinning Disk Confocal Microscopy

For synaptome mapping, we used Spinning Disk confocal Microscopy (SDM) platforms (Figure 2). The Andor Revolution XDi was used with an Olympus UPlanSAPO 100X oil immersion lens (NA 1.4), a CSU-X1 spinning-disk (Yokogawa) and an Andor iXon Ultra monochrome back-illuminated EMCCD camera. Images acquired with this system have a pixel dimension of 84 × 84 nm and a depth of 16 bits. A single mosaic grid was used to cover each entire brain section with an adaptive Z focus set-up by the user to follow the unevenness of the tissue using Andor iQ2 software. The field of view of each individual frame was 43.008 x 43.008 μm. In both systems, eGFP was excited using a 488 nm laser and mKO2 with a 561 nm laser. The CV1000 system is equipped with the following filters: BP 525/50 nm for eGFP and BP 617/73 nm for mKO2, whereas the Andor Revolution XDi is equipped with a Quad filter (BP 440/40, BP 521/21, BP 607/34 and BP 700/45). For both systems, mosaic imaging was set up with no overlap between adjacent tiles.

**Figure 2.**
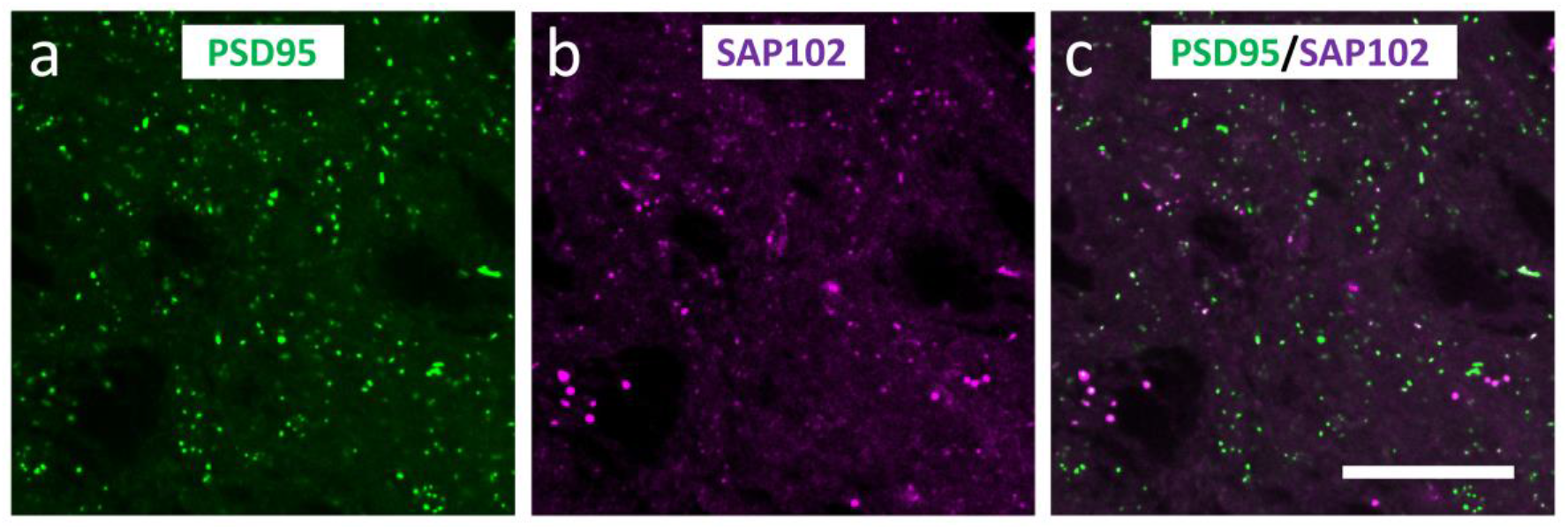
Spinning Disk confocal Microscopy (SDM). **(a** and **b)**: Examples of PSD95 fluorescent puncta (green) and SAP102 puncta (purple), imaged with SDM in the stratum lacunosum-moleculare of CA1. The PSD95 and SAP102 channels have been merged in **(c)**. Calibration bar: 15 μm.

### Detection and measurement of fluorescent Synaptic Puncta

Punctum detection was performed using Ensemble Detection, an in-house collection of image detection algorithms. We have developed a new punctum/particle detection method based on a multi-resolution image feature detector and supervised machine learning technique (Zhu et al., 2018). In this method, we carry out a multi-resolution and multi-orientation version of 2nd-order nonlocal derivative (NLD) (Qiu et al., 2012), and use it to calculate intensity differences, referred to as ‘image features’, for each of the individual puncta at different spatial resolutions and orientations. An initial intensity threshold is set to a very low value to only filter out extremely dim puncta and to avoid missing true synaptic puncta. The remaining candidate puncta were finally classified as either true puncta or background noise using the corresponding feature vectors and the classifier. The classifier was pre-trained with the training image set and machine learning algorithms (Qiu et al., 2012).

After detection and localization of all puncta, we segmented them based on their individual intensity values: for each punctum, a threshold was set as 10% of the maximum pixel intensity within the punctum, so that punctum size and shape measurement were independent of punctum intensity (Zhu et al., 2018). With the puncta segmented and binarized, six punctum parameters were then calculated: mean punctum pixel intensity, punctum size, skewness, kurtosis, circularity, and aspect ratio.

### Tissue Preparation for electron microscopy

Four male PSD95^eGFP/eGFP^; SAP102^mKO2/Y^ mice were used for electron microscopy. Animals were administered a lethal intraperitoneal injection of sodium pentobarbital (40 mg/kg) and were intracardially perfused with 2% paraformaldehyde and 2.5% glutaraldehyde in 0.1 M phosphate buffer (PB). The brain was then extracted from the skull and processed for EM as previously described (Merchán-Pérez et al., 2009). Briefly, the brains were post-fixed at 4°C overnight in the same solution used for perfusion. They were then washed in PB and vibratome sections (150 μm thick) were obtained. Sections containing the rostral hippocampus were selected with the help of an atlas (Paxinos and Franklin, 2004). Selected sections were osmicated for 1 hour at room temperature in PB with 1% OsO4, 7% glucose and 0.02 M CaCl2. After washing in PB, the sections were stained for 30 min with 1% uranyl acetate in 50% ethanol at 37°C, and they were then dehydrated and flat embedded in Araldite (DeFelipe and Fairén, 1993). Embedded sections were glued onto blank Araldite stubs and trimmed. To select the exact location of the samples, we first obtained semithin sections (1–2 μm thick) from the block surface and stained them with toluidine blue to identify cortical layers. These sections were then photographed with a light microscope. The last of these light microscope images (corresponding to the section immediately adjacent to the block face) was then collated with low power scanning electron microscope (SEM) photographs of the surface of the block. In this way, it was possible to accurately identify the three strata of the hippocampus to be studied.

### Three-Dimensional Electron Microscopy

Three-dimensional brain tissue samples of the CA1 of the hippocampus were obtained using combined focused ion beam milling and scanning electron microscopy (FIB-SEM) (Figure 3). The focus of our study was the neuropil, which is composed of axons, dendrites and glial processes. We used a CrossBeam 540 electron microscope (Carl Zeiss NTS GmbH, Oberkochen, Germany). This instrument combines a high-resolution field emission SEM column with a focused gallium ion beam, which can mill the sample surface, removing thin layers of material on a nanometer scale. After removing each slice (20 nm thick), the milling process was paused, and the freshly exposed surface was imaged with a 1.8-kV acceleration potential using the in-column energy selective backscattered (EsB) electron detector. The milling and imaging processes were sequentially repeated, and long series of images were acquired through a fully automated procedure, thus obtaining a stack of images that represented a three-dimensional sample of the tissue (Merchán-Pérez et al., 2009). Twelve samples (stacks of images) of the neuropil of three strata of CA1 were obtained, avoiding the neuronal and glial somata as well as the blood vessels (Figure 4). These stacks included four samples of stratum lacunosum moleculare (SLM), four of stratum radiatum (SR) and four of stratum oriens (SO) (see Supplementary Table 1). In these stacks, we obtained the densities of glutamatergic (asymmetric) and GABAergic (symmetric) synaptic junctions. To do this, we counted the number of synaptic junctions within an unbiased three-dimensional counting frame of known volume (Howard and Reed, 2005). Image resolution in the xy plane was 5 nm/pixel; resolution in the z-axis (section thickness) was 20 nm and image sizes were 2048 x 1536 pixels (field of view: 10.24 x 7.68 μm). The number of sections per stack ranged from 201 to 377 (mean 276.33; total 3316 sections). Processing for EM causes shrinkage of the tissue for which we have to correct the measurements (Merchán-Pérez et al., 2009). Correction factors for the tissue that was used in theis study were 0.9508 for linear measurements, 0.9040 for area measurements and 0.8595 for volumetric data. The volumes of the stacks, after correction for tissue shrinkage, ranged from 367.81 to 689.86 μm^3^ (mean 505.66 μm^3^; total 6067.86 μm^3^). The volumes of the counting frames ranged from 288.62 to 585.99 μm^3^ (mean 408.76 μm^3^; total 4905.07 μm^3^) (Supplementary Table 1).

**Figure 3.**
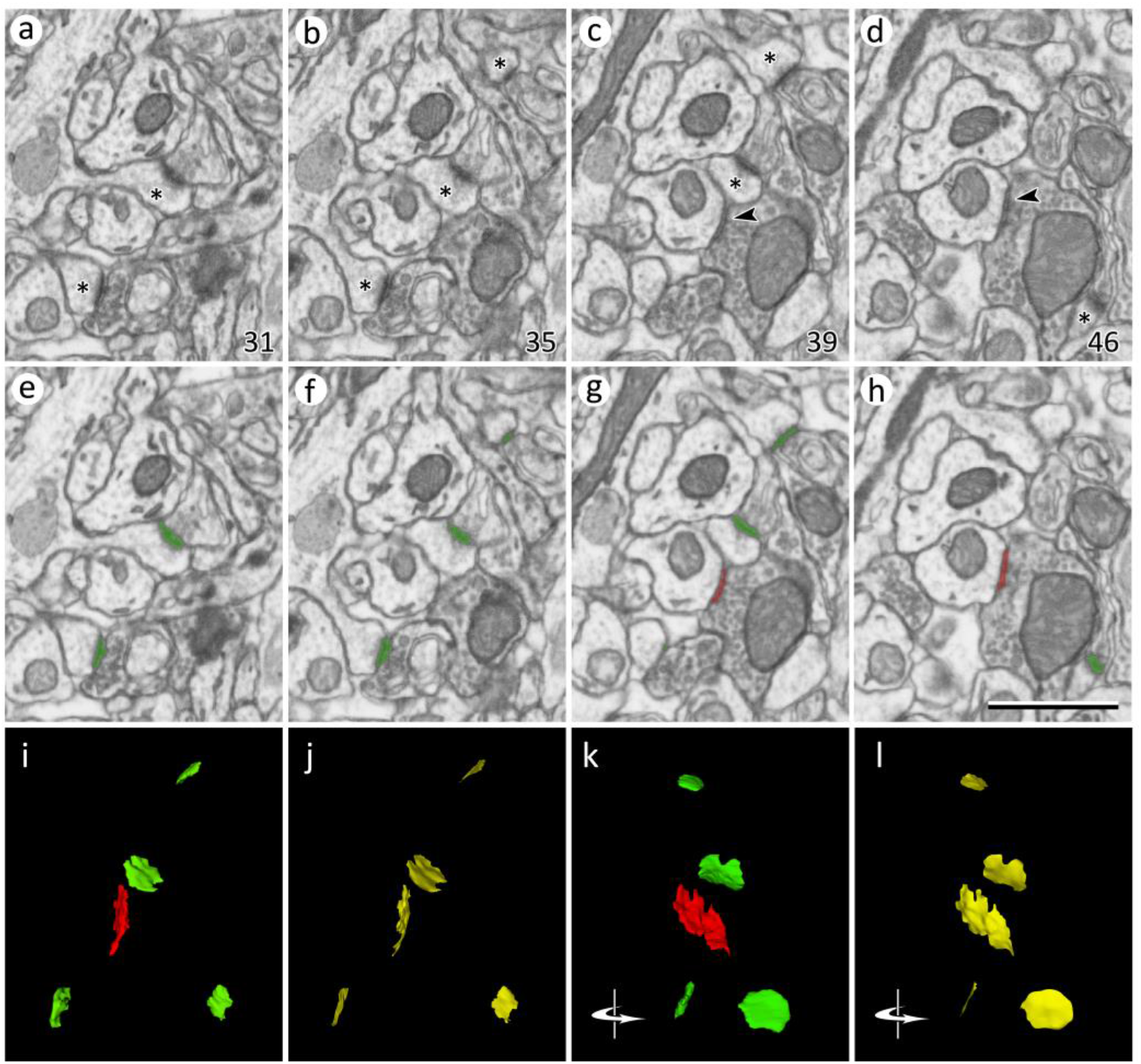
Identification and segmentation of synaptic junctions in serial sections acquired by FIB-SEM. **(a-c)** Detail of four electron micrographs selected from a series o images obtained by FIB-SEM. In this example, the stack of images was obtained from the stratum oriens. The numbers in the bottom-right corner correspond to section number. Four asymmetric synapses can be identified by the presence of prominent post-synaptic densities in (a), (b), (c) and (d) (asterisks). One symmetric synapse, with a thin post-synaptic density, can be seen in (c) and (d) (arrow heads). Note that the classification of synapses as asymmetric or symmetric is not based on single images but on the examination of the full sequence of images. **(e-h)** The same images after they have been segmented with Espina software (http://cajalbbp.es/espina/). The segmentation process is based on grey-level thresholds, so the resulting 3D objects comprise both the pre- and post-synaptic densities (see methods). Green profiles correspond to asymmetric synapses and red profiles to the symmetric synapse. **(i)** 3D rendering of the synaptic junctions present in (a) to (h). **(j)** Synaptic apposition surfaces (SAS, yellow) extracted from the 3D segmentations represented in (i). SAS are automatically extracted from the 3D reconstructions of synaptic junctions (see methods); they are zero-volume surfaces that represent the interface between the pre- and post-synaptic densities. The surface area of the SAS is measured for each individual synaptic junction. **(k, l)** Same structures represented in (i) and (j), respectively, after they have been rotated through a vertical axis. Original images were acquired with a resolution of 5 nm/pixel, with a distance of 20 nm between two consecutive images. Calibration bar in (h) is 1 μm.

**Figure 4.**
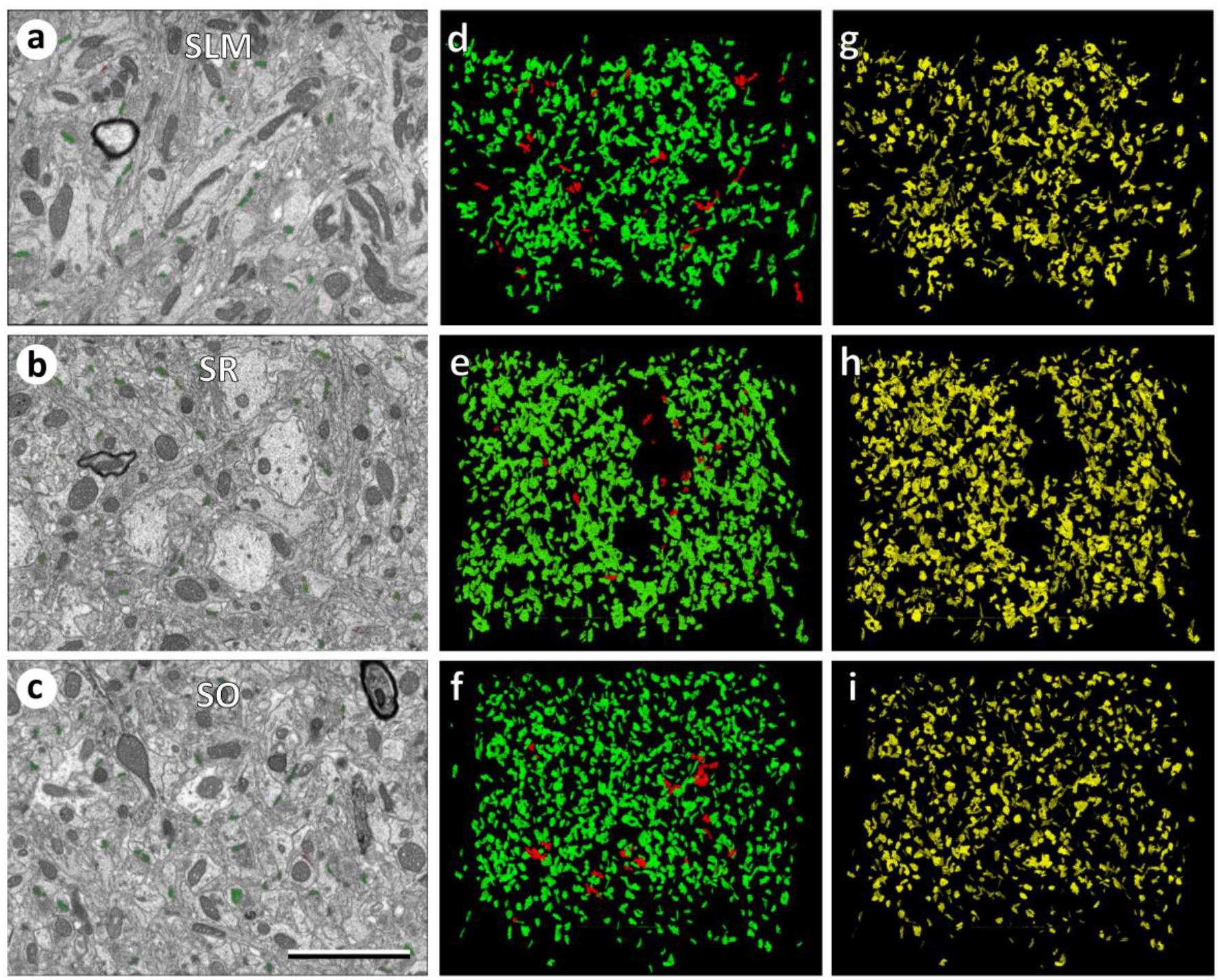
Measuring synaptic densities and sizes in stacks of sections obtained by FIB-SEM. **(a-c)** Panoramic view of electron micrographs of the stratum lacunosum moleculare (SLM), stratum radiatum (SR) and stratum oriens (SO) imaged by FIB-SEM. **(d-f)** 3D rendering of synaptic junctions reconstructed from the corresponding stacks of serial sections, acquired from the strata represented in (a) to (c). Asymmetric synaptic junctions have been represented in green and symmetric synaptic junctions in red. **(g-i)**. The synaptic apposition surfaces (SAS, yellow) have been automatically extracted from the three-dimensionally reconstructed synaptic junctions. The number of synapses per unit volume and the surface areas of the SAS have been measured in each stack of serial sections (see Tables 1 and 2). Calibration bar in (c): 3 μm.

### Identification and reconstruction of synapses

Synaptic junctions within these volumes were visualized and segmented in 3D with Espina software (Morales et al., 2011) (http://cajalbbp.es/espina/). The segmentation algorithm makes use of the fact that presynaptic and postsynaptic densities appear as dark, electron-dense structures under the electron microscope. It requires a Gaussian blur filter preprocessing to eliminate noisy pixels and then it uses a gray-level threshold to extract all the voxels that fit the gray levels of the synaptic junction. In this way, the resulting 3D segmentation includes both the active zone (AZ) and postsynaptic density (PSD) (Morales et al., 2013). Synaptic junctions with a prominent or thin PSD were classified as asymmetric or symmetric synaptic junctions, respectively (Colonnier, 1968; Gray, 1959) (Figure 3). Synapses could be unambiguously identified since they can be visualized in consecutive serial sections and, if necessary, they can be digitally resectioned in different planes to ascertain their identity as asymmetric or symmetric synapses (DeFelipe et al., 1999; Merchán-Pérez et al., 2009).

### Size of synapses

As stated above, the synaptic junction is formed by the AZ and the PSD. Since AZ and PSD are in close apposition and have similar surface areas, they can be represented as a single surface — the synaptic apposition surface (SAS). Thus, the SAS is an accurate measurement of the size of the synapse. In previous studies we have developed an efficient computational technique to automatically extract this surface from reconstructed synapses (Morales et al., 2013).

### Statistical analysis

To study whether there were significant differences between synaptic distributions among the different CA1 layers, we performed a multiple mean comparison test. When the data met the criteria of normality and homoscedasticity, an ANOVA was performed. When these criteria were not met, we used the Kruskal-Wallis followed by Dunn’s test for pair-wise comparisons.

## Results

We estimated the density and the size of synapses in the CA1 area of the hippocampus using two different methods (Figure 1). PSD95-positive and SAP102-positive synapses were identified as fluorescent puncta using SDM (Figure 2), and FIB-SEM was used to visualize and reconstruct synaptic junctions in the same regions. FIB-SEM also provided information that was not obtained from confocal images, such as the relative proportions of excitatory (asymmetric) and inhibitory (symmetric) synapses (Figure 3, Supplementary Table 1). This classification of synapses is based on the appearance of the PSD in EM images (Colonnier, 1968; Gray, 1959). Any synaptic junction with a dense, prominent PSD that was much thicker than the relatively faint presynaptic thickening was classified as “asymmetric” (AS). Any synapse with a less marked PSD, similar to the presynaptic thickening, was classified as “symmetric” (SS) (Merchán-Pérez et al., 2009). It should be stressed that the classification of synaptic junctions into one of these two groups was not based on the examination of single sections, but on the whole series of images in which the PSD was visible (Figure 3). Once all synapses within a given stack of serial sections had been identified and segmented, they appeared as a cloud of 3D objects from which quantitative data were obtained (Figure 4).

### Density of fluorescent puncta and synapses in the hippocampus

Densities of fluorescent puncta (number of positive puncta per 100 μm^2^) were measured in SLM, SR and SO from CA1. Sixteen brain sections were used (one section per animal, see Supplementary Table 2). We obtained the densities of puncta expressing PSD95 *(dPSD95)* and SAP102 *(dSAP102),* as well as the colocalization index (*c*). From these data we calculated the total density of puncta *(dTotal)* (Table 1, Supplementary Table 2). Note that *dTotal* is not simply the sum of dPSD95 and dSAP102, since there is a certain density of puncta that colocalize (*dColoc*):

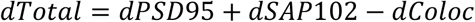

**Table 1.** Relationship between the densities of puncta and the densities of synapses. PSD95 and SAP102 puncta were imaged with SDM. The total densities of puncta were calculated from the densities of PSD95 and SAP102 puncta, together with the colocalization index (see text for details). The densities of asymmetric synapses (AS) were obtained from volumes of tissue reconstructed from serial sections using FIB-SEM. For each layer, the conversion factors is the quotient between the density of synapses obtained by FIB-SEM and the total density of puncta calculated from SDM images. Densities are given as average ± SD

| Stratum of CA1 | Density of PSD95 (puncta/100 $\mu\text{m}^2$) | Density of SAP102 (puncta/100 $\mu\text{m}^2$) | Colocalization index | Total density of puncta/100 $\mu\text{m}^2$ | Density of AS (synapses/$\mu\text{m}^3$) | Conversion factor |
| --- | --- | --- | --- | --- | --- | --- |
| Lacunosum-Moleculare | 91.3103 ± 13.3605 | 77.1788 ± 23.8497 | 0.5840 | 106.4372 ± 17.6450 | 1.5958 ± 0.7317 | 0.0150 |
| Radiatum | 118.4513 ± 25.8022 | 122.7022 ± 24.2477 | 0.6633 | 145.1760 ± 25.6250 | 2.3076 ± 0.3788 | 0.0159 |
| Oriens | 119.7759 ± 21.1004 | 115.6872 ± 18.9181 | 0.6741 | 140.7479 ± 17.2418 | 2.4887 ± 0.4763 | 0.0177 |
| All layers averaged | 109.8458 ± 24.2337 | 105.1894 ± 29.8734 | 0.6404 | 130.7871 ± 26.6441 | 2.1307 ± 0.6396 | 0.0162 |

The colocalization index (*c*) ranges from 0, when there is no colocalization, to 1, when there is 100% colocalization, so *dColoc* is related to *dTotal* according to the following expression:

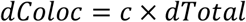

Therefore,

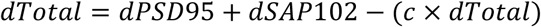

from which we obtain:

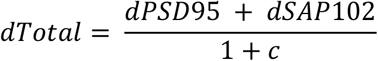

We observed the highest total density of puncta in SR (mean ± SD; 145.18 ± 25.62 puncta/100 μm^2^), followed by SO (140.75 ± 17.24 puncta/100 μm^2^) and SLM (106.44 ± 17.64 puncta/100 μm^2^) (Table 1, Supplementary Table 2). The differences between SLM and the other two layers were statistically significant (KW test, p < 0.005) (Figure 5a).

**Figure 5.**
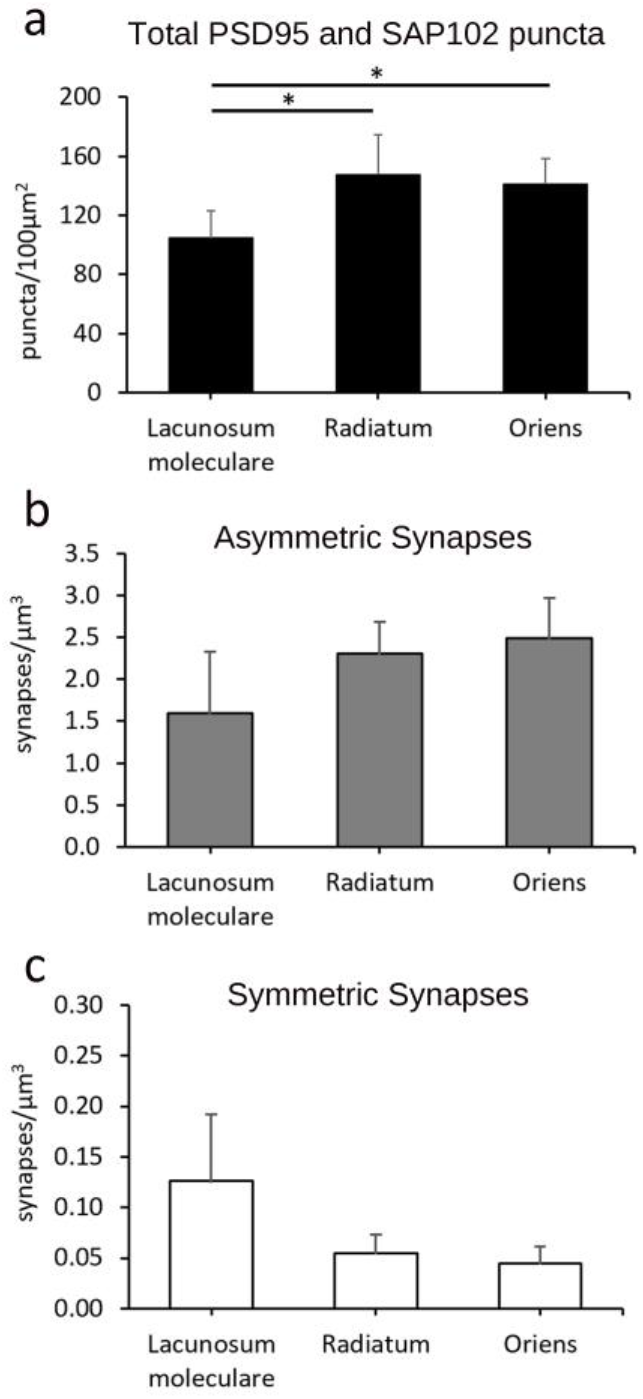
Total densities of PSD95 and SAP102 puncta, asymmetric synapses and symmetric synapses. (**a**) Density of PSD95- and SAP102-positive puncta (puncta/100 μm^2^ ± SD) acquired by SDM in the hippocampus (CA1). Asterisks indicate statistically significant differences (KW, p < 0.005). (**b)** and **(c)** Density of asymmetric and symmetric synapses, respectively (synapses/μm^3^ ± SD), estimated from stacks of serial sections acquired by FIB-SEM from the same regions. See also Table 1.

For the volume electron microscopy study (FIB.SEM), we used 12 stacks of serial sections from SLM, SR and SO (Figure 3, Figure 4, Supplementary Table 1). In these samples, we identified and analyzed a total of 10,460 synapses in 4,905 μm^3^ of tissue. Of these, 95.60% were AS and 4.40% were SS. To estimate the density of synapses in each stack of images, we counted the number of synaptic junctions within an unbiased three-dimensional counting frame of known volume (see Methods). The density of AS (mean ± SD) in SLM was 1.59 ± 0.73 synapses/μm^3^, in SR it was 2.31 ± 0.38 synapses/μm^3^, and in SO it was 2.49 ± 0.48 synapses/μm^3^ (Table 1).

SS were most frequent in SLM (0.13 ± 0.07 synapses/μm^3^), followed by SR (0.06 ± 0.02 synapses/μm^3^) and SO (0.05 ± 0.02 synapses/μm^3^). In spite of this trend of an increase in AS density from SLM to SO, and a decrease in SS across these strata, the differences between layers were not statistically significant for either the total density of synapses (AS+SS) or for AS and SS separately (KW test, p ≥ 0.08) (Figure 5b, c).

### Size of fluorescent puncta and synapses in the hippocampus

We measured the area of PSD95 puncta in SLM, SR and SO. The largest mean area of puncta was found in SLM (0.0832 μm^2^), followed by SR (0.0809 μm^2^) and SO (0.0798 μm^2^) (Figure 6a). Although the differences were small, they were statistically significant (KW test p < 0.001).

**Figure 6.**
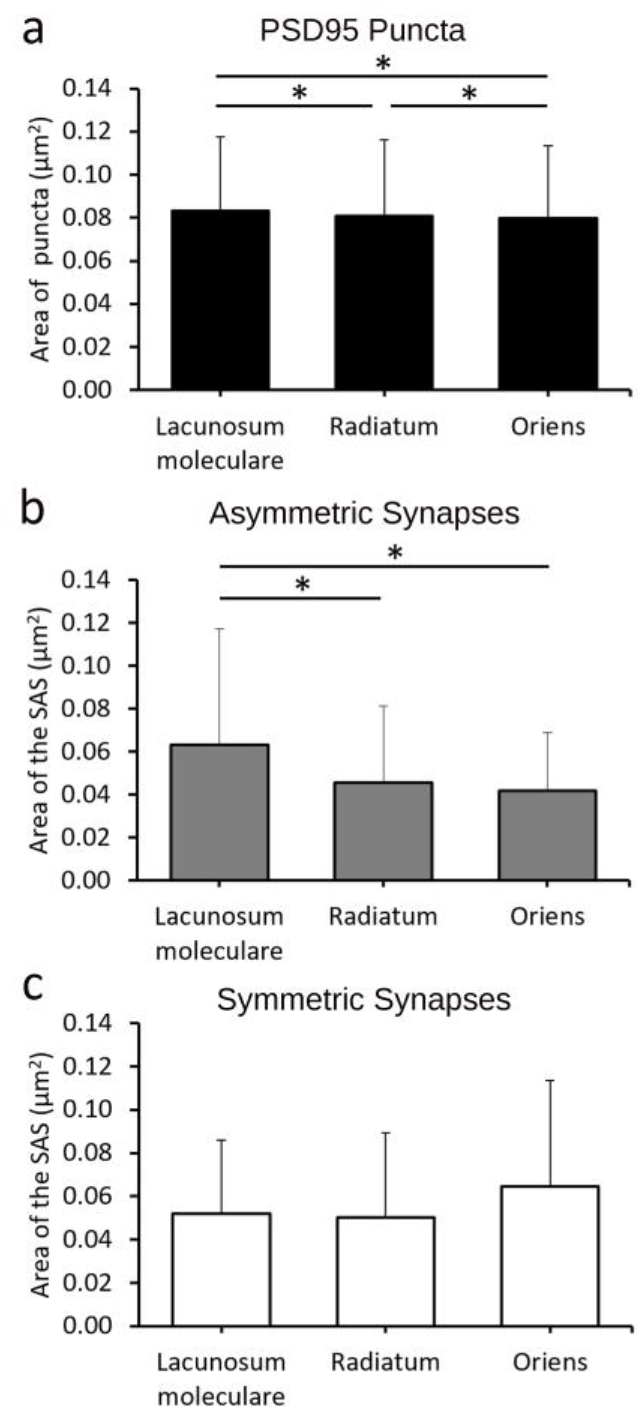
Size of PSD95 puncta, asymmetric synapses and symmetric synapses. **(a)** Area of PSD95-positive puncta (μm^2^) acquired by SDM in the hippocampus (CA1) (Mean + SD). Asterisks indicate statistically significant differences. **(b)** and **(c)** Mean size of the synaptic apposition surface (SAS) of asymmetric and symmetric synapses (μm^2^) estimated from stacks of serial sections acquired by FIB-SEM from the same region. See also Table 2.

To estimate the size of synapses in the FIB-SEM samples, we measured the area of the synaptic apposition surface (SAS). The SAS is a surface that represents the apposition between the presynaptic density and the PSD and reproduces their curvature (Figure 3, Figure 4, see Methods). The mean SAS area in the FIB-SEM samples was 0.0474 μm^2^ for asymmetric synapses and 0.0541 μm^2^ for symmetric synapses.

For asymmetric synapses, the mean SAS area in SLM was larger than in the other layers (0.0633 μm^2^; KW-Dunn’s p < 0.001)(Figure 6b). Despite the mean SAS area being larger in SR than in SO (0.0456 μm^2^ and 0.0419 μm^2^, respectively), the difference was not statistically significant (KW-Dunn’s p > 0.05). For symmetric synapses, the largest mean SAS areas were found in SO (0.0644 μm^2^) followed by SLM (0.0519 μm^2^) and SR (0.0501 μm^2^) (KW, p = 0.05) (Figure 6c, Table 2).

**Table 2.** Surface areas of PSD95 puncta acquired by SDM, and surface areas of the synaptic apposition surface (SAS) of asymmetric (AS) and symmetric (SS) synapses reconstructed from FIB-SEM samples. The number of puncta or synapses analyzed (n), as well as the parameters μ and σ of the corresponding best-fit log-normal distributions are also indicated.

|  |  | Stratum |  |  |  |
| --- | --- | --- | --- | --- | --- |
|  |  | Lacunosum-Moleculare | Radiatum | Oriens | All layers |
| PSD 95 Puncta | Mean Area ± SD ($\mu\text{m}^2$) | $0.0832 \pm 0.0342$ | $0.0809 \pm 0.0352$ | $0.0798 \pm 0.0336$ | $0.0813 \pm 0.0342$ |
|  | n | 147776 | 189945 | 125916 | 463637 |
| | $\mu$ | 11.24 | 11.20 | 11.20 | 11.21 |
| | $\delta$ | 0.44 | 0.48 | 0.46 | 0.46 |
| AS | Mean SAS Area ± SD ($\mu\text{m}^2$) | $0.0633 \pm 0.0540$ | $0.0456 \pm 0.0358$ | $0.0419 \pm 0.0271$ | $0.0474 \pm 0.0377$ |
|  | n | 2258 | 4538 | 5082 | 11878 |
| | $\mu$ | 10.74 | 10.55 | 10.50 | 10.54 |
| | $\delta$ | 0.80 | 0.63 | 0.57 | 0.64 |
| SS | Mean SAS Area ± SD ($\mu\text{m}^2$) | $0.0520 \pm 0.0338$ | $0.0501 \pm 0.0394$ | $0.0644 \pm 0.0491$ | $0.0541 \pm 0.389$ |
|  | n | 247 | 93 | 87 | 427 |
| | $\mu$ | 10.67 | 10.60 | 10.97 | 10.70 |
| | $\delta$ | 0.68 | 0.65 | 0.75 | 0.69 |

When we compared the sizes of asymmetric synapses and symmetric synapses in different layers, we found that symmetric synapses were larger than asymmetric synapses in SO and SR, while in SLM the opposite was the case. The greatest differences were found in SO, where mean SAS areas for symmetric synapses and asymmetric synapses were in a proportion of approximately 6:4 (MW test, p < 0.0001) (Table 2).

To further characterize the size distribution of synaptic sizes, we plotted the frequency histograms of the areas of PSD95 puncta and of the SAS (Figure 7). The frequency histograms of the areas of PSD95 puncta showed skewed shapes, with a long tail to the right. The histograms of SAS areas of asymmetric synapses measured from FIB-SEM reconstructions also showed skewed shapes, but they were narrower and lay to the left of PSD95 histograms in all layers (Figure 7).

**Figure 7.**
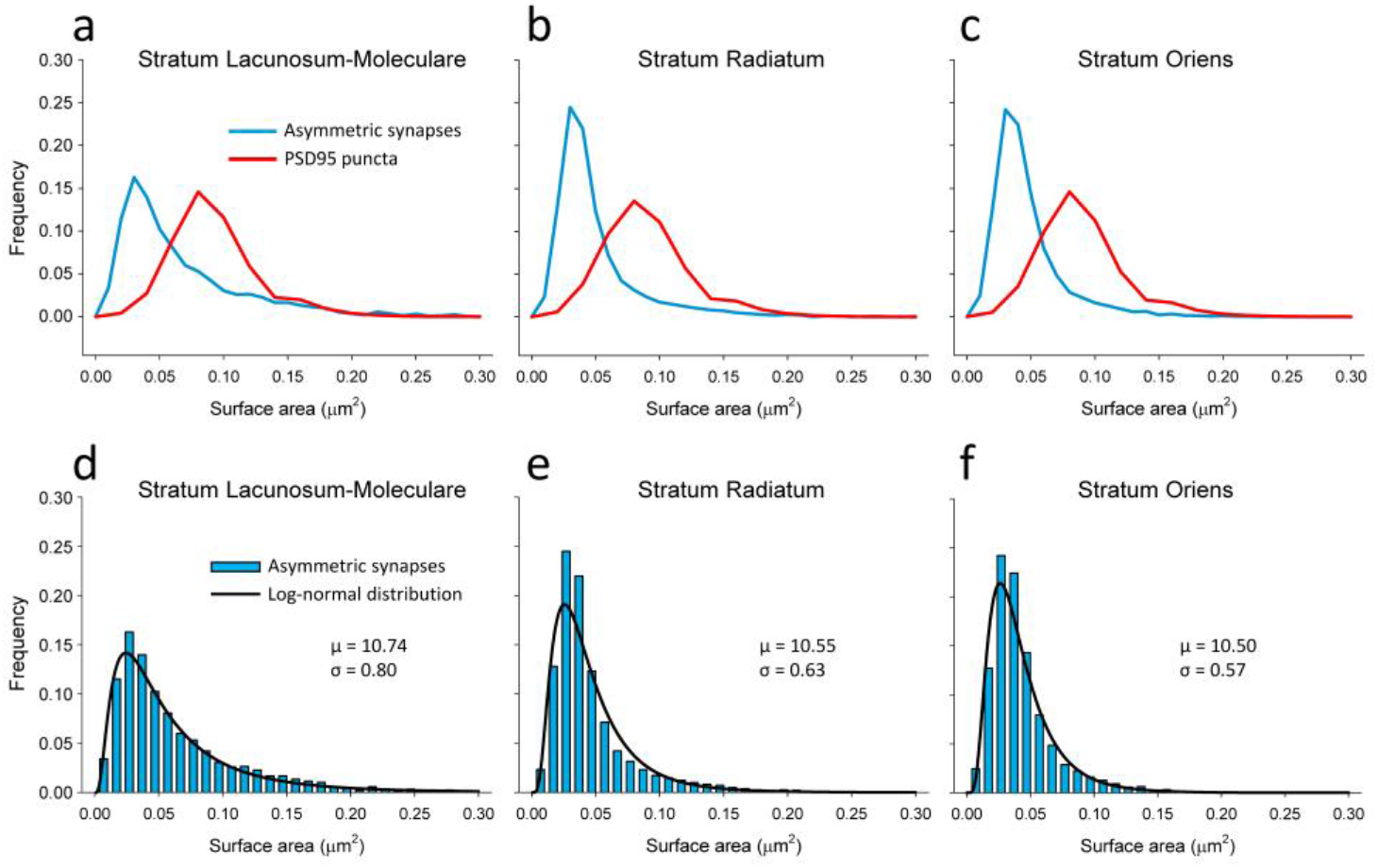
Frequency histograms of the sizes of PSD95 puncta and asymmetric synapses. **(a-c)** Comparison of the distribution of the surface areas of PSD95 puncta acquired by SDM (red line) and the synaptic apposition surfaces of asymmetric synapses reconstructed from FIB-SEM samples (blue line) from three layers of the CA1 region of the hippocampus. The histograms corresponding to asymmetric synapses are narrower and lie to the left of the histograms corresponding to PSD95 puncta. **(d-f)** Frequency histograms (blue bars) of the areas of the synaptic apposition surfaces of asymmetric synapses reconstructed from FIB-SEM samples. The log-normal distributions (black lines) represent the theoretical probability density functions that best fit the experimental data. The parameters μ and σ of the corresponding log-normal distributions have also been indicated.

We then performed goodness-of-fit tests to find the theoretical probability density functions that best fitted the empirical distributions of the areas of PSD95 puncta and SAS areas. We found that they fitted to log-normal distributions in all cases, with some variations in the parameters μ and σ (Table 2 and Figure 7).

### Brain-wide estimations of the number of synapses

When we compared the densities of PSD95 and SAP102 puncta measured with SDM and the densities of AS measured with FIB-SEM, we found that both methods revealed that SR and SO had similar densities, while SLM had a lower density (Figure 5). We then calculated a conversion factor that would allow us to relate the densities of PSD95 and SAP102 puncta (puncta/100 μm^2^) to the actual densities of excitatory synapses found by FIB-SEM (synapses/μm^3^). These conversion factors were calculated as the quotient between the actual density of AS and the total density of PSD95 and SAP102 puncta. Conversion factors obtained for each layer of CA1 were slightly different; they ranged from 0.0152 to 0.0176. The averaged conversion factor calculated with data from the three layers was 0.0162 (Table 1).

The next step was to calculate the total number of puncta expressing PSD95 and/or SAP102 brain-wide, using previously published data from 113 areas (Zhu et al., 2018). Different brain regions had different combinations of densities of PSD95, SAP102 and total densities of puncta (Figure 8, Supplementary Table 2). The highest total densities of puncta were found in the isocortex, the olfactory areas, the hippocampal formation and the cortical subplate. All these regions were relatively homogeneous except for the hippocampal formation, which showed wider ranges of variability. Also, isocortical areas had a relatively higher proportion of PSD95 versus SAP102 than the other regions. The pallidum, the hypothalamus, the brainstem and the cerebellum had low densities of puncta. Finally, the striatum —and especially the thalamus— showed the greatest variability. For example, of all thalamic nuclei, the ventral medial nucleus had one of the lowest estimated densities of total puncta (19.78 puncta/100 μm^2^), while the posterior complex had one of the highest estimated densities (110.15 puncta/100 μm^2^) (Supplementary Table 2). As a validation step, we compared previously published data regarding CA1 (Zhu et al., 2018) with our present data; the total densities of puncta expressing PSD95 and/or SAP102 were remarkably similar (128.95 puncta/100μm^2^ in previously published data and 130.96 puncta/100μm^2^ in the present study).

**Figure 8.**
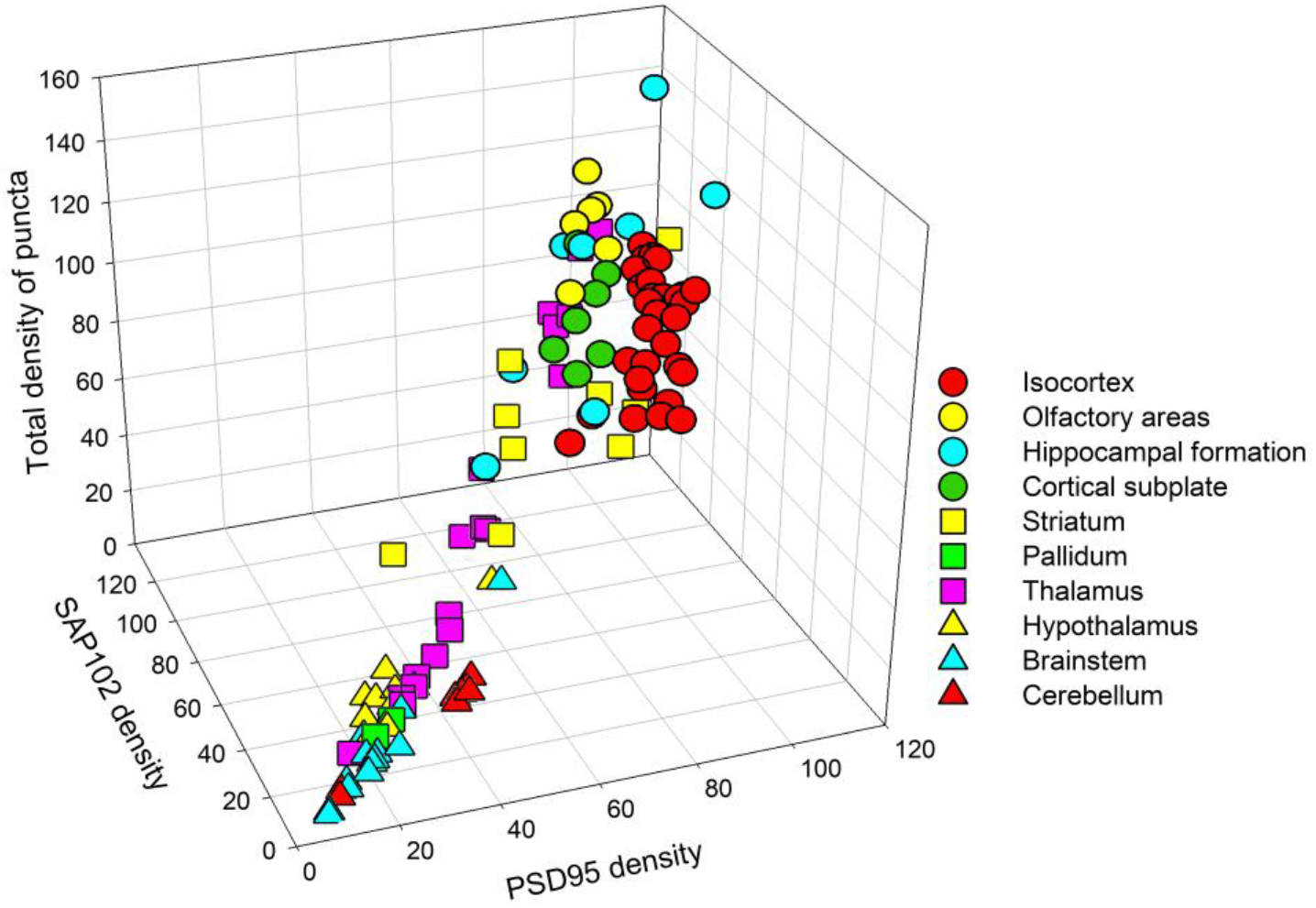
Densities of PSD95 and SAP102 puncta, and total densities of puncta in different regions of the brain. The total densities of puncta have been calculated from previously published densities and colocalization indexes of PSD95 and SAP102 puncta (Zhu et al., 2018). Symbols represent the different subregions within the major brain regions listed in the legend (see Supplementary Table 2 for the complete list of subregions). The isocortex, the olfactory areas, the hippocampal formation and the cortical subplate have high densities of puncta. The pallidum, the hypothalamus, the brainstem and the cerebellum have low densities of puncta. There is a wide variability in the total densities of puncta in the striatum and thalamus.

Finally, we estimated the density of synapses expressing PSD95 and/or SAP102, making use of the averaged conversion factor obtained in CA1 (Table 1). The values obtained have been graphically represented in Figure 9. In general, the hippocampal cornu ammonis, the isocortex and the olfactory areas had the highest synaptic densities, intermingled with cortical subplate nuclei. Within the hippocampal formation, the dentate gyrus and the subiculum presented similar densities, but these were lower than in the Ammon’s horn. Striatal nuclei showed considerable variations, but the thalamic nuclei showed the highest variability, as mentioned above. The cerebellar cortex showed homogeneously low densities and the pallidum, hypothalamus and brainstem had the lowest synaptic densities.

**Figure 9.**
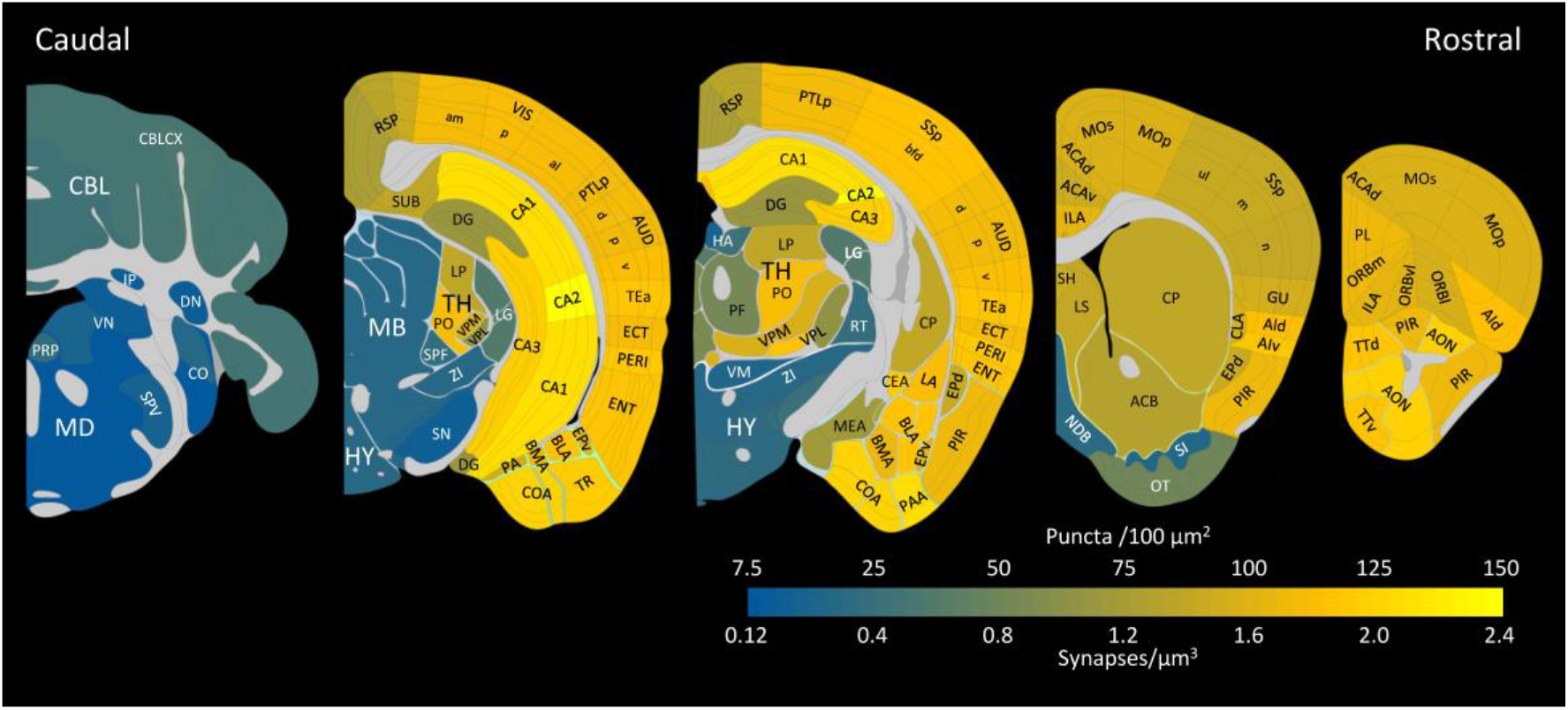
Estimated densities of synapses expressing PSD95 and/or SAP102 in different regions of the brain. The total densities of puncta per 100 square microns have been calculated from previously published densities of PSD95 and SAP102, as well as their colocalization indexes (Zhu et al., 2018). A conversion factor obtained in CA1 was used to estimate the minimum densities of synapses per cubic micron (see text for details). Illustrations and brain regions are based on the Allen Mouse Brain Atlas. The following structures have been labeled: Cerebellum (CBL), CBLCX: cerebellar cortex; DN: Dentate nucleus; IP: Interposed nucleus. Medulla (MD), CO: Cochlear nuclei; PRP: Nucleus prepositus; SPV: Spinal nucleus of the trigeminal; VN: Vestibular nuclei. Midbrain (MB), SN: Substantia nigra. Hypothalamus (HY), ZI: Zona incerta. Thalamus (TH), HA: Habenular nuclei; LG: Lateral geniculate complex; LP: Lateral posterior nucleus; PF: Parafascicular nucleus; PO: Posterior complex; RT: Reticular nucleus; SPF: Suprafascicular nucleus; VM: Ventral medial nucleus; VPL: Ventral posterolateral nucleus; VPM: Ventral posteromedial nucleus. Pallidum, NDB: Diagonal band nucleus; SI: Substantia innominata. Striatum, ACB: Nucleus accumbens; CEA: Central amygdalar nucleus; CP: Caudoputamen; LS: Lateral septal nucleus; MEA: Medial amygdalar nucleus. OT: Olfactory tubercle; SH: septohippocampal nucleus. Cortical subplate, BLA: Basolateral amygdalar nucleus; BMA: Basomedial amygdalar nucleus; CLA: Claustrum; EPd: Endopiriform nucleus, dorsal part; EPv: Endopiriform nucleus, ventral part; LA: Lateral amygdalar nucleus; PA: Posterior amygdalar nucleus. Olfactory areas, AON: Anterior olfactory nucleus; COA: Cortical amygdalar area; PAA: Piriform-amygdalar area; PIR: Piriform area; TR: Postpiriform transition area; TTd: Taenia tecta, dorsal part; TTv: Taenia tecta, ventral part. Hippocampal formation, CA1, CA2, CA3: Cornu Ammonis, fields 1, 2 and 3; DG: Dentate gyrus. ENT: Entorhinal area; SUB: Subiculum. Isocortex, ACAd: Anterior cingulate area, dorsal part; ACAv: Anterior cingulate area, ventral part; AId: Agranular insular area, dorsal part; AIv: Agranular insular area, ventral part; AUD: Auditory areas (d, p, v: dorsal, primary ventral); ECT: Ectorhinal area; GU: Gustatory area; ILA: Infralimbic area; MOp: Primary motor area; MOs: Secondary motor area; ORBl: Orbital area, lateral part; ORBm: Orbital area, medial part; ORBvl: Orbital area, ventrolateral part; PERI: Perirhinal area; PL: Prelimbic area; PTLp: Posterior parietal association areas; RSP: Retrosplenial area; SSp: Primary somatosensory area (bfd, ul m, and n: barrel field, upper limb, mouth and nose representations); TEa: Temporal association areas; VIS: Visual areas (am, p, al: anteromedial, primary, anterolateral).

## Discussion

In this study, we have —for the first time— analyzed the synaptic density of excitatory and inhibitory synapses, as well as their size, in stratum oriens, stratum radiatum and stratum lacunosum-moleculare of the CA1 hippocampal region of the mouse, using three-dimensional electron microscopy. With this method, long series of consecutive sections are obtained by FIB-SEM, so individual synapses can be unambiguously identified and the number of synapses per unit volume can be directly calculated. However, as with any other electron-microscopy technique, FIB-SEM can only be applied to relatively small regions of tissue, so it is not practical for brain-wide estimations. By contrast, the number of fluorescent puncta expressing PSD95 and/or SAP102 can be quantified brain-wide using SDM, so we have attempted to establish a correlation between the two kinds of measurements.

### Synaptic sizes and densities in the hippocampus

Regarding the size of fluorescent puncta and PSDs, what is actually measured with SDM imaging is different to what is measured after reconstruction of synaptic junctions from serial images obtained by FIB-SEM. In the case of SDM, the images obtained are two-dimensional, so what we actually see is the two-dimensional projection of puncta on the plane of section. Thus, puncta with different orientations will show different apparent surface areas, and only those that are oriented flat with respect to the plane of section will show their true surface area. By contrast, serial images obtained by FIB-SEM allow us to reconstruct the synaptic junctions in 3D. We can then extract the synaptic apposition surface (SAS) from each individual synapse. The SAS represents the surface of apposition between the presynaptic and postsynaptic densities, so the surface area of the SAS is equivalent to the area of the PSD, and we can measure it for every synapse, regardless of its spatial orientation (Morales et al., 2013).

We have found that SDM imaging clearly overestimates the size of PSD95 puncta when compared with the actual size of PSDs imaged by FIB-SEM (see Figure 7). This can be due to several factors. Light scatter, glare and blur may contribute to the fact that fluorescent puncta appear to be larger than the actual PSDs. The resolution of SDM is also much lower than the resolution of electron microscopy. In the x-y plane, the resolution of SDM was 84 nm/pixel, while FIB-SEM images were acquired at a resolution of 5 nm/pixel. This makes a pixel area of 7056 nm^2^ for SDM versus only 25 nm^2^ for FIB-SEM. The lower resolution may result in SDM missing the smaller synapses and those that are oriented perpendicularly to the plane of section. Also, the images of several synapses may overlap throughout the thickness of the SDM optical section. As a result, some puncta may in fact be clusters of two or more synapses. In spite of these differences between the two-dimensional SDM imaging and volume electron microscopy, the measurements of fluorescent puncta by SDM do distinguish the relative size differences between layers or regions, so they are still useful for the identification and classification of synaptic types (Zhu et al., 2018). Both our SDM and FIB-SEM results indicate that excitatory synapses in SLM are larger than in the SR or SO, in line with previous studies in the rat (Megías et al., 2001).

The distribution of synaptic sizes measured from FIB-SEM stacks of images fits a log-normal distribution in the three strata analyzed (see Figure 7d-f). This trait has also been described in the rat neocortex (Merchán-Pérez et al., 2014; Santuy et al., 2018b). This type of distribution is characterized by a skewed curve with a long tail to the right, and it has been found in other synaptic parameters such as synaptic strength, spike transmission probability, and the size of unitary excitatory postsynaptic potentials (Buzsáki and Mizuseki, 2014; Lefort et al., 2009; Song et al., 2005; Hazan and Ziv, 2020). It is thus tempting to suggest that the size of the synaptic junction is correlated with these and other functional characteristics of the synapse, as has been proposed previously (Santuy et al., 2018b).

Regarding the densities of puncta and synapses in the hippocampus, previous studies in the rat SR reported 2.2 synapses/μm^3^ using EM and three-dimensional reconstructions (Mishchenko et al., 2010), and similar estimates using stereological methods (Sorra et al., 1998). In both cases, the reported synapse densities were lower than the density we have found in the mouse SR (2.4 synapses/μm^3^). Differences between species may explain the discrepancies, although we cannot rule out the possibility of other sources of bias, such as the different methods used. On the other hand, our SDM results regarding the total density of PSD95 and SAP102 puncta in the hippocampus were very similar to previously reported data (Zhu et al., 2018). We also provide information about the amount of inhibitory synapses, represented by symmetric or type 2 synapses (Colonnier, 1968; Gray, 1959). These do not express PSD95 or SAP102, so their densities cannot be estimated from our SDM data. However they can be identified in FIB-SEM images because of their thin PSD (Merchán-Pérez et al., 2009). In our CA1 samples, symmetric synapses represented 4.4% of the total number of synapses. This is in line with results in SR and SO of the rat CA1, where percentages of inhibitory synapses as low as 3% have been reported in thin dendrites, which predominate in our samples (Megías et al., 2001). Interestingly, they also reported that —in line with our results— SLM had the highest percentage of inhibitory synapses (leaving aside the stratum pyramidale and the thick proximal dendrites, which were not included in our study).

### Brain-wide estimations of the minimum densities of synapses

We next applied a conversion factor obtained in the hippocampus to calculate synaptic densities brain-wide. The conversion factor was calculated as the ratio between the densities of excitatory (asymmetric) synapses obtained by FIB-SEM and the total density of PSD95 and SAP102 puncta obtained by SDM. We found that the conversion factors were very similar in the three CA1 layers studied, and we used an averaged conversion factor for brain-wide estimations (see Table 1). It is important to bear in mind the limitations of this procedure to ensure that the results are interpreted correctly.

While it is clear that only excitatory glutamatergic synapses express PSD95 and/or SAP102 (Nithianantharajah et al., 2013; Frank et al., 2017; Zhu et al., 2018), the question of whether *all* excitatory synapses express these scaffolding proteins does not have a simple answer. In the adult mouse hippocampus, it has been recently claimed that all Schaffer collateral/commissural synapses in the SR of CA1 show immunogold staining for PSD95 (Yamasaki et al., 2016). This is probably an overestimate, since our own data indicate that there is a population of synapses that do not express PSD95, but do express SAP102 (Supplementary Table 2). In any case, if we consider that Schaffer collateral/commissural fibers are the origin of the vast majority of synapses in SR and SO, we can assume that most, if not all, synapses in these strata express PSD95, SAP102 or a combination of the two. It is likely to be the same case in SLM, since the ratio between the number of fluorescent puncta and the actual density of synapses measured by FIB-SEM is very similar to that of the two other layers (see Table 1). Lower percentages of immunolabeling of synapses with PSD95 have been reported in the rat hippocampus (Sans et al., 2000), but this has been attributed to the low sensitivity of the technique (Sassoé-Pognetto et al., 2003). In our case, the advantage of the genetic labeling method is that all PSD95 and SAP102 proteins are labeled, so a more reliable detection is to be expected.

However, even if we assume that the vast majority of excitatory synapses in CA1 express PSD95 and/or SAP102, and that we can detect them in a reliable way, the question remains as to whether this would be the case in other brain regions. For example, PSD95 was regarded as “a fundamental structural component of most, if not all, excitatory PSDs isolated from the rat cerebral cortex” (Petersen et al., 2003). Other studies seem to confirm this view (Swulius et al., 2010; DeGiorgis et al., 2006), while lower percentages of PSD95-expressing synapses have also been reported (Aoki et al., 2001; Farley et al., 2015). Brain-wide studies in the mouse seem to confirm that the abundance of scaffolding proteins like PSD95 and SAP102 differs depending on the brain area (Roy et al., 2018; Zhu et al., 2018). Also, although PSD95 and SAP102 together are thought to label most excitatory synapses, we expect that labeling with PSD93 could reveal an additional set of excitatory synapses (Elias et al., 2006). Finally, the vast majority of PSD95 puncta in the adult brain are found in the postsynaptic terminals, but they have also been observed in non-synaptic locations in developing neurons (Gerrow et al., 2006; Washbourne et al., 2002). Thus, we cannot rule out the possibility that some of the PSD95 puncta observed in the adult mouse brain are not from extra-synaptic sites.

Therefore, to interpret our results correctly, we must clearly assume that not all excitatory synapses throughout the brain express PSD95 and/or SAP102. Since our calculations are based only on the population of synapses that express these scaffolding proteins, our estimations of synaptic densities do in fact underestimate the actual densities of excitatory synapses. In other words, our estimations represent the lower boundary of the densities of excitatory synapses in different brain regions. The upper boundary cannot be estimated from our present data, since this would require knowing the proportion of excitatory synapses that *do not* express PSD95 and/or SAP102 in each brain region.

Only a systematic exploration of the different regions of the brain with FIB-SEM or similar methods will settle the possible discrepancies between our present estimations and the actual values. However, we can compare our estimations with previous studies, when available. In the mouse neocortex, previously reported synaptic densities using different stereological methods were either lower (Sadaka et al., 2003; Schüz and Palm, 1989) or higher (DeFelipe et al., 1997) than our present estimations. In the juvenile rat somatosensory cortex, the mean density of synapses in the neuropil has been reported to be between 0.87 and 0.89 synapses/μm^3^ using FIB-SEM (Anton-Sanchez et al., 2014; Santuy et al., 2018a), which is below our present estimation for the adult mouse somatosensory cortex (1.4 to 1.9 synapses/μm^3^, see Supplementary Table 2). However, these differences may be due to species and/or age differences (e.g., DeFelipe et al., 1997). In the rat cerebellum, the density of synapses has been previously reported to be 0.8 synapses/μm^3^ in the molecular layer (Napper and Harvey, 1988), while our present estimations for the mouse cerebellar cortex range from 0.5 to 0.6 synapses/μm^3^ (Supplementary Table 2). Therefore, we currently lack data that are directly comparable to our present estimations, since methodological bias is probably at play in those cases, leaving aside the possible species and age differences. Although work is already in progress on the mouse somatosensory cortex using a FIB-SEM methodology that is similar to the one presented here, it would not be practical to wait until results from even a fraction of the 113 subregions examined here become available. Therefore, our calculations must be regarded as reasonable —but provisional— estimations of the minimum densities of glutamatergic synapses in the different brain regions.

In summary, it is important to emphasize that acquiring multiple samples at different scales is a highly effective way to obtain a dataset that allows comprehensive analysis of the brain. Since the whole brain cannot be fully reconstructed at the ultrastructural level, it seems clear that only by combining studies at the meso- and nano-scopic levels (light and electron microscopy) can we fully understand the structural arrangement of the brain as a whole [see, for example (Markram et al., 2015, Kashiwagi et al., 2019)]. Using this strategy, we provide an estimation of the minimum densities of glutamatergic synapses in the different brain regions. These data, in combination with previous studies on the relationship between the connectome and synaptome (Zhu et al., 2018), can be used to identify common and differing principles of synaptic organization. This in turn could serve to further advance efforts to validate and refine realistic brain models.

## Acknowledgements

This work was supported by grants from the following entities: the Spanish “Ministerio de Ciencia, Innovación y Universidades” (grant PGC2018-094307-B-100 and the Cajal Blue Brain Project [C080020-09; the Spanish partner of the Blue Brain Project initiative from EPFL, Switzerland]; the European Union’s Horizon 2020 Research and Innovation Programme under grant agreement No. 785907 (Human Brain Project, SGA2); the Wellcome Trust (Technology Development Grant 202932); and the European Research Council (ERC) under the European Union’s Horizon 2020 research and innovation programme (695568 SYNNOVATE). L.T.-R. is a recipient of grants from the EMBO Long-term fellowship 2016–2018 and the IBRO-PERC InEurope grants programme.

## Notes

### Competing Interest Statement

The authors have declared no competing interest.

