## Supplementary material for "Estimation of the number of synapses in the hippocampus and brain-wide by volume electron microscopy and genetic labeling": Suppl Table 1

| Stratum | Sample ID | Animal ID | No. of serial sections | Section thickness (nm) | Resolution (nm/pixel) | Total volume ($\mu m^3$) | CF volume ($\mu m^3$) | No. of AS | No. of SS | No. of synapses | Density of AS (synapses / $\mu m^3$) | Density of SS (synapses / $\mu m^3$) | Density of AS+SS (synapses / $\mu m^3$) |
|---|---|---|---|---|---|---|---|---|---|---|---|---|---|
| Lacunosum moleculare | 1 | PSD95-ID7 | 273 | 20 | 5 | 499.5554 | 337.2320 | 341 | 69 | 410 | 1.0112 | 0.2046 | 1.2158 |
| | 8 | PSD95-ID10 | 295 | 20 | 5 | 539.8126 | 427.6239 | 569 | 67 | 636 | 1.3306 | 0.1567 | 1.4873 |
| | 15 | PSD95-ID15 | 212 | 20 | 5 | 387.9331 | 303.2372 | 417 | 24 | 441 | 1.3752 | 0.0791 | 1.4543 |
| | 18 | PSD95-ID16 | 230 | 20 | 5 | 420.8708 | 316.1865 | 843 | 20 | 863 | 2.6661 | 0.0633 | 2.7294 |
| Radiatum | 17 | PSD95-ID15 | 240 | 20 | 5 | 439.1695 | 364.7681 | 1026 | 26 | 1052 | 2.8127 | 0.0713 | 2.8840 |
| | 20 | PSD95-ID16 | 240 | 20 | 5 | 439.1695 | 344.0458 | 689 | 23 | 712 | 2.0026 | 0.0669 | 2.0695 |
| | 3 | PSD95-ID7 | 312 | 20 | 5 | 570.9204 | 474.6386 | 964 | 15 | 979 | 2.0310 | 0.0316 | 2.0626 |
| | 10 | PSD95-ID10 | 344 | 20 | 5 | 629.4763 | 505.8370 | 1206 | 25 | 1231 | 2.3842 | 0.0494 | 2.4336 |
| Oriens | 2 | PSD95-ID7 | 307 | 20 | 5 | 561.7710 | 432.0351 | 1082 | 14 | 1096 | 2.5044 | 0.0324 | 2.5368 |
| | 11 | PSD95-ID10 | 377 | 20 | 5 | 689.8622 | 585.9860 | 1587 | 24 | 1611 | 2.7083 | 0.0410 | 2.7492 |
| | 16 | PSD95-ID15 | 201 | 20 | 5 | 367.8045 | 288.6178 | 843 | 20 | 863 | 2.9208 | 0.0693 | 2.9901 |
| | 19 | PSD95-ID16 | 285 | 20 | 5 | 521.5138 | 524.8589 | 956 | 19 | 975 | 1.8214 | 0.0362 | 1.8576 |
| Totals | Lacunosum moleculare | | | | | | 1384.2796 | 2170 | 180 | 2350 | 1.5958 | 0.1259 | 1.7217 |
| | Radiatum | | | | | | 1689.2895 | 3885 | 89 | 3974 | 2.3076 | 0.0548 | 2.3624 |
| | Oriens | | | | | | 1831.4979 | 4468 | 77 | 4545 | 2.4887 | 0.0447 | 2.5334 |
| | | | | | | | 4905.0670 | 10523 | 346 | 10869 | 2.1307 | 0.0751 | 2.2059 |

**Supplementary Table 1.** Stacks of serial sections obtained by FIB SEM and used for the estimation of densities of synapses in the CA1 region of the hippocampus.
CF: counting frame. AS: asymmetric synapses. SS: symmetric synapses.
